# Rapid, precise quantification of large DNA excisions and inversions by ddPCR

**DOI:** 10.1101/2020.04.13.039297

**Authors:** Hannah L. Watry, Carissa M. Feliciano, Katie Gjoni, Gou Takahashi, Yuichiro Miyaoka, Bruce R. Conklin, Luke M. Judge

## Abstract

The excision of genomic sequences using paired CRISPR-Cas nucleases is a powerful tool to study gene function, create disease models and holds promise for therapeutic gene editing. However, our understanding of the factors that favor efficient excision is limited by the lack of a rapid, accurate measurement of DNA excision outcomes that is free of amplification bias. Here, we introduce ddXR (<u>d</u>roplet <u>d</u>igital PCR e<u>X</u>cision <u>R</u>eporter), a method that enables the accurate and sensitive detection of excisions and inversions independent of length. The method can be completed in a few hours without the need for next-generation sequencing. The ddXR method uncovered unexpectedly high rates of large (>20 kb) excisions and inversions, while also revealing a surprisingly low dependence on linear distance, up to 170 kb. We further modified the method to measure precise repair of excision junctions and allele-specific excision, with important implications for disease modeling and therapeutic gene editing.

## Introduction

Gene editing using CRISPR-Cas9 or other nuclease systems in eukaryotic cells occurs via precisely targeted double-stranded DNA cleavage events, followed by repair by endogenous cellular machinery^1^. The Cas9 nuclease finds its target via a guide RNA (gRNA) with complementarity to the desired locus^2^. Many gene editing approaches rely on imprecise non-homologous end-joining (NHEJ) repair to create small insertions and deletions (indels) that disrupt a gene at a single cut site. Other approaches rely on homology-directed repair (HDR) to introduce a new sequence at the double-stranded break. Larger editing events can be produced by simultaneously delivering two nucleases targeted to different sequences on the same locus, which can lead to large deletions via excision of the intervening genomic sequence^3–7^. Generating excisions with paired CRISPR gRNAs is an attractive means to engineer complete loss of gene function, map regulatory regions, study 3D genome organization and model deletion-induced diseases. Furthermore, various therapeutic applications utilize paired gRNA to remove precise regions of DNA to induce alternate exon splicing, inactivate dominant disease alleles, remove toxic repeat expansions and delete viral integrations^8–15^. However, dual gRNA editing produces multiple editing outcomes, including excisions and inversions between the two cut sites, and indels at one or both sites^3,5^. There is currently no reliable approach to predict the relative frequency of these outcomes, or to measure it accurately and efficiently.

Small indels produced at individual target sites are routinely assessed via sequencing PCR amplicons by either Sanger or next generation sequencing (NGS), including by our group^16–20^. However, the quantification of excisions by amplicon sequencing or other measurements of amplicon abundance is complicated by amplification bias due to the inherent size difference between the amplicons of the excised and non-excised alleles, limiting this approach to very short excisions^8,16,17^. Detection of inversions requires additional specific primers, adding further complexity and risk of differential amplification of multiple amplicons even if they are designed to be similar in size. The extensive optimization required to validate every combination of primers for each editing experiment makes this an impractical approach. Unfortunately, the primary alternative approach to quantifying large excision events has been to isolate large numbers of cell clones for genotyping and sequencing each clone^3,5^. This process is slow and labor intensive, with sensitivity limited by the number of clones that can be analyzed. It is also limited to proliferative cell lines that can undergo clonal isolation. Whole-genome sequencing (WGS) has been used to measure the frequency of a small excision, but is expensive, low throughput and read depth limits its ability to detect infrequent events^21^. Targeted single molecule DNA sequencing could provide a useful alternative, but remains expensive, may suffer from differential selection of excised sequences and be limited by maximum read length^22,23^. UDiTaS, a unidirectional sequencing method, is able to quantify excisions and inversions in population samples without amplification bias. However, UDiTaS requires additional investment, including novel library preparation, NGS and downstream computational analysis^24^. None of these methods allow for rapid, low cost, reliable and length-independent quantification of excisions at endogenous loci in a heterogeneous population.

Here, we introduce ddXR (<u>d</u>roplet <u>d</u>igital PCR e<u>X</u>cision <u>R</u>eporter) to enable the sensitive and precise quantification of excisions and inversions without apparent length limitations. Droplet digital PCR (ddPCR) has several advantages over standard PCR for the detection of gene editing events. It is highly sensitive and quantitative^25^ and has previously been used to measure NHEJ and HDR editing outcomes^26–28^. Furthermore, by encapsulating target DNA molecules in individual droplets before amplification, it minimizes the problem of amplification bias. Finally, data can be analyzed immediately without downstream library preparation or computational expertise. The ddXR protocol can be completed the same day as DNA extraction and produces a gain of signal (GOS) that makes it possible to measure even rare excision and inversion events in mixed edited populations. We demonstrate the accuracy and consistency of this assay to detect excisions and inversions ranging from 91 bp to 172 kb in length. We also describe further modifications of the method to measure precise repair events and to measure allele-specific excision in a model of dominant genetic disease. The speed, simplicity and versatility of ddXR make it an ideal standard for the quantification of excisions and inversions in genome editing experiments.

## Results

### ddXR accurately detects excisions and inversions over broad length and frequency ranges

We first tested our ddXR approach with pairs of Cas9/gRNA ribonucleoprotein (RNP) complexes targeting two genomic loci present at two copies in the reference human genome. We designed assays to produce a gain of signal measurable by ddPCR for excision and inversion occurring at each target locus (Fig. 1a,b). For excisions, we designed primers and FAM-labelled probes flanking the nuclease target sites so that they would be brought into close proximity after excision occurs, allowing for efficient amplification and activation of the fluorophore (Fig. 1a,b). We reasoned that unedited alleles would not produce a signal, as amplification of the intact sequence would be inefficient. For inversions, the same FAM probe and its associated primer were used, along with an alternate second primer located between the gRNA target sites and targeting the same strand as the first primer. Inversion reverses the orientation of the second primer and brings it into proximity with the first primer and probe, allowing for specific amplification (Fig. 1a). Combining this design with digital PCR technology allows amplification to occur at the single-molecule level, critical for minimizing amplification bias. Finally, we added an internal control assay for a reference gene, RPP30, also present at two copies in the human genome. The RPP30-specific probe was labelled with HEX, which allowed us to easily calculate the proportion of alleles with an excision or inversion in our target gene by normalizing the FAM signal to the HEX signal.

**Figure 1:**
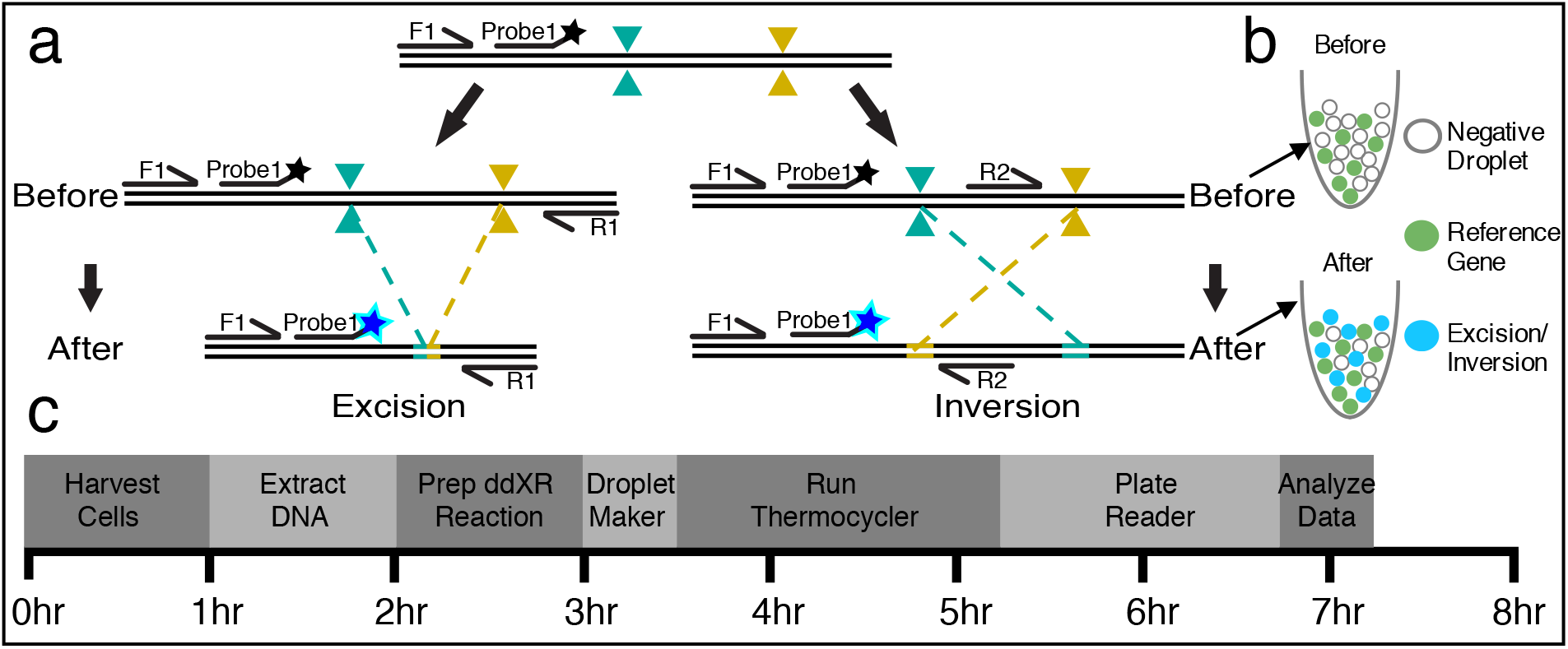
Overview of ddXR excision and inversion quantification methods. A) Schematic of ddXR excision and inversion assay designs. Both inversion and excision occur in the same sample, but they are assayed in separate reactions. Note that the only variation between the excision and inversion assay is the reverse primer. Forward (F1) and reverse primers (R1, R2) are indicated by arrows along with FAM-conjugated fluorescent probe. Primers and probes to the reference gene are not shown. B) Illustration of expected droplets detected before (upper) and after (lower) editing is performed. The number of excision or inversion positive droplets (blue) is normalized to the number of RPP30 positive droplets (green). RPP30 is a reference gene present at two copies in the human genome. C) Workflow of DNA extraction and ddXR assay with time estimates.

To validate the method, we transfected euploid, human iPSCs with the two pairs of RNP complexes. One pair was located 4.09 kb apart on chromosome 8 and the other 1.7 kb apart on chromosome 7. We isolated clones with heterozygous and homozygous deletion of 4.09 kb on chromosome 8 and heterozygous inversion of 1.7 kb on chromosome 7, as determined by traditional PCR (Fig. 2a,b). Based on the FAM-to-HEX ratio, we obtained modest levels of excision (12.5%) and inversion (4.8%) in the edited iPSC populations. Clonal iPSC lines produced expected ratios of excised and inverted alleles—48.3% and 98.7% in heterozygous and homozygous excision lines, respectively, and 50.2% in a heterozygous inversion line (Fig. 2c,d). We isolated additional clonal excision lines, with excisions on chromosomes 7 and 8, and again detected heterozygous excisions with the expected frequency (53.5% and 47.8%, respectively; Fig. S1). Next, we asked whether locating the probe at the 5’ or 3’ end of an excision would affect the results of the assay. We tested this in the clone with the largest excision (14kb on chromosome 1) and found no significant difference in excision frequency measured by the two probes (Fig. S2). We did not observe false positive signals from unedited control samples in any of these assays, although we did detect a high false positive rate when generating very short excisions (<200bp). This was caused by efficient amplification of the non-excised alleles and could be easily removed by an internal restriction digest (Fig. S3).

**Figure 2:**
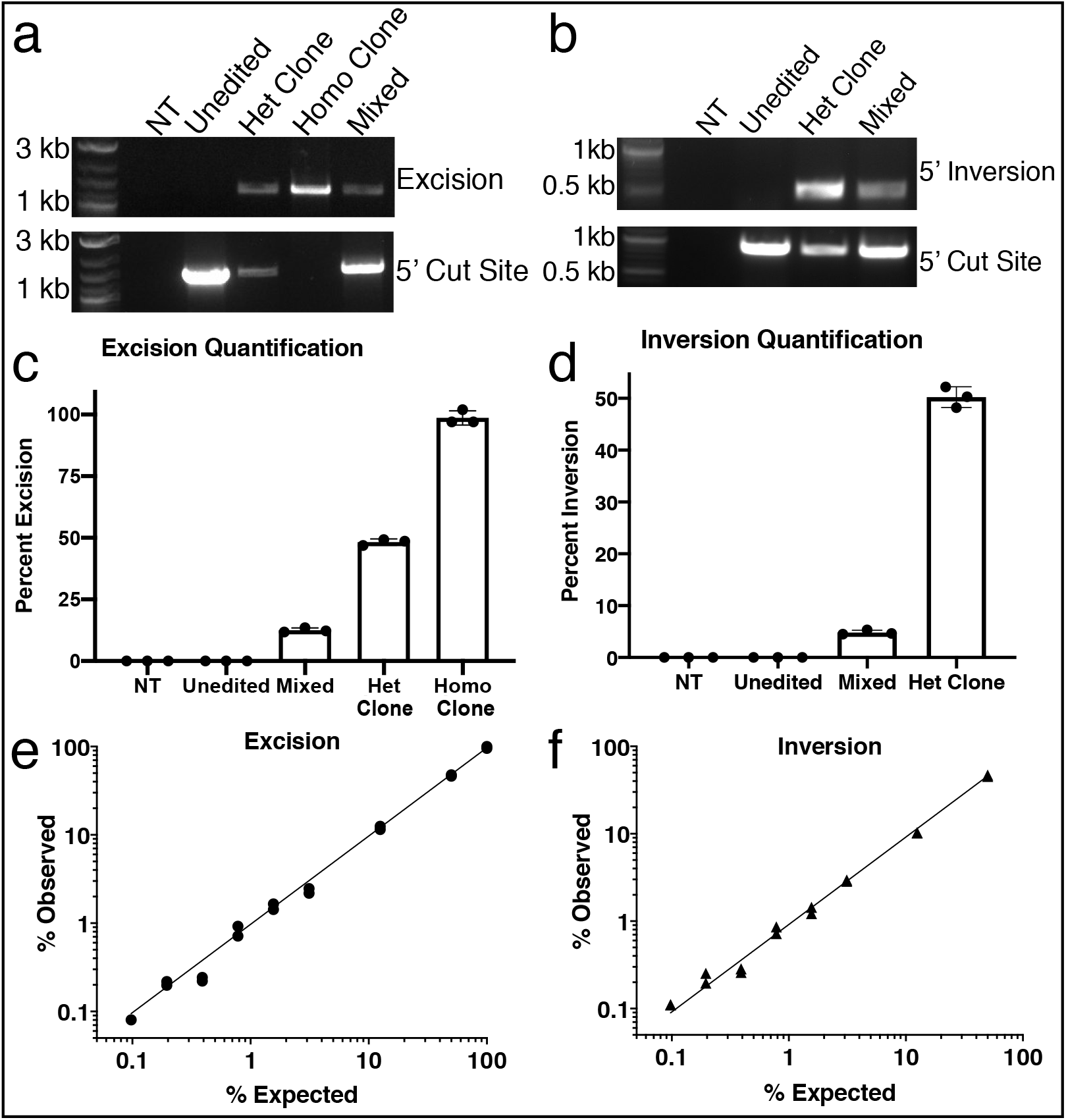
Validation of ddXR using clonal iPSC lines to measure excision and inversion. A,B) Qualitative detection of excision/inversion and characterization of clonal cell lines used to validate assay. Edited clones were derived from previously characterized, karyotypically normal iPSC lines. PCR assays specific for a 4kb excision on chromosome 8 (A) or 1.7kb inversion on chromosome 7 (B) produced bands in mixed samples and clones. NT= no template. Control PCR spanning 5’ cut site detects alleles without excision/inversion and produced bands in the unedited sample, heterozygous clones and mixed sample. We did not identify any clones with homozygous inversion. C,D) Quantification of excision (C) and inversion (D) in mixed and clonal populations using ddXR assay. Values represent the percent of total alleles with excision/inversion and are averages of three replicates. E,F) ddXR quantification of standard samples with defined frequency of excision (E) and inversion (F). Samples prepared by serial dilution of clonal DNA with the unedited parental line. Data include two independent replicates of each dilution and are presented as the percentage of alleles with excision/inversion detected by ddXR. Note that in (F) the maximum inversion frequency is 50% as it was prepared using genomic DNA from a heterozygous clone. R^2^ = 0.999 (E) and 0.998 (F).

To evaluate the dynamic range of ddXR, we generated genomic DNA samples with known proportions of either excised or inverted alleles. We prepared two independent serial dilutions of genomic DNA from two sets of edited clones (homozygous excision and heterozygous inversion clones) with genomic DNA from the unedited parental clone. We tested each dilution series with the ddXR assay and obtained results that were highly accurate, even when measuring samples with excision or inversion frequencies as low as 0.1% (Fig. 2e,f). A previously reported approach for quantifying excision frequency with ddPCR used the loss of signal (LOS) of an internal probe^29^. We repeated the analysis of the standardized excision samples using a LOS ddPCR assay. In this case, there was reduced accuracy and linearity, with root mean square error over four times higher than that for the corresponding ddXR assay (Fig. S4). The LOS method performed particularly poorly at lower concentrations of excision alleles.

We next compared the frequency of excisions and inversions measured by ddXR to those determined by PCR genotyping of individual iPSC clones. We handpicked 48 colonies and genotyped the surviving clones from four distinct paired RNP transfections (Table S1). We then compared the proportion of excision alleles based on clone genotyping to the ddXR quantification from the same transfection. There was reasonable agreement between the methods, with one experiment producing a modest discrepancy between ddXR and clone genotyping (6.8% vs. 11.5% respectively, Table S1). However, there was variability in the number of surviving clones between experiments, with low total numbers of positive clones limiting the statistical power of this approach. These results provided independent confirmation of the excision and inversion rates measured by ddXR, while highlighting the enhanced speed, precision and cost effectiveness of ddXR.

### Excision and inversion correlate poorly with length

The factors that influence excision efficiency are not well understood, although it is widely believed that there is a strong negative correlation with increasing linear distance between cut sites^3^. We applied ddXR to investigate this possibility in three genes that differ in size, numbers of exons, chromosome and expression patterns. We designed gRNAs and ddXR assays and transfected human iPSCs with various pairs of RNP complexes targeting each locus, with a linear distance between target sites ranging from 91 bp to 172 kb (Tables S2,3). Genomic DNA was isolated from each transfected population and assayed for both excision and inversion (Fig. 3a,b, S5). Interestingly, we saw wide variation in excision and inversion frequencies across the full range of linear distance between cut sites, with only a weak correlation between excision and linear distance and no correlation between inversion and linear distance (Fig. 3.a,b). We additionally observed a significantly lower overall rate of inversions compared to excisions (Fig. 3c). The longest excision (172kb on chromosome 1) occurred at an unexpectedly high frequency (9.87%, Fig. 3a), presenting the opportunity to validate this result by fluorescent in-situ hybridization (FISH). FISH was performed using one probe targeting the deleted fragment and a second control probe targeting the same chromosome 5.5 Mb away. Chromosomes containing the excision were identified by positive signal for the control probe but negative signal for the target probe, which occurred in 10% of alleles in the transfected population (Fig. S6a,c). This was highly consistent with the excision frequency of 9.87% determined by ddXR on the same sample (Fig. S6a). No chromosomes showed loss of target signal in the unedited control by FISH (Fig S6b).

**Figure 3:**
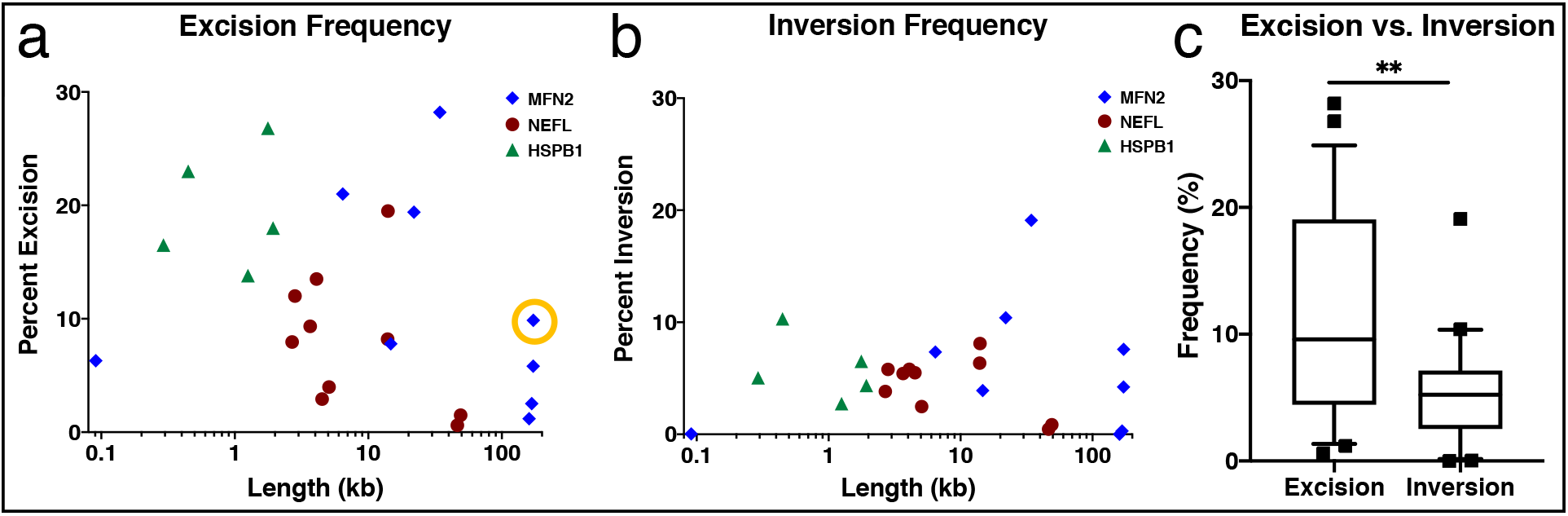
Quantification of excision and inversion frequencies versus linear distance between cut sites. A,B) Dot plots of all excisions (A) and inversions (B) measured at three loci with varying linear distance between paired CRISPR-Cas9 target sites. Color and shape of points indicate locus. Pearson correlation suggests that excision is negatively correlated with increasing length, but with only borderline significance. r = −0.409, p = 0.047 (A) and detected no correlation for inversions, r = −0.216, p = 0.310 (B). Circled excision rate in (A) was validated by FISH (Fig. S6). C) Box plot comparing overall frequency of excisions and inversions across the entire dataset as percentage of total alleles. Whiskers extend to 10^th^ and 90^th^ percentile. Outliers are marked with black squares. Median_excision_ = 9.589%, Median_inversion_ = 5.230%, p = 0.0015.

### ddXR distinguishes between precise and imprecise excisions

When sequencing excision junctions, we noticed that many samples had small insertions or deletions at the repair junction, consistent with other types of NHEJ repair. The ability to produce precise excision events is an important tool for studies of biological function as well as gene therapy^6,15^. To determine the frequency of these events, we designed an assay to detect precise excision repair, where the two cut sites are ligated together without additional insertions or deletions. This assay uses the same primers as the general ddXR excision quantification assay, but a probe that spans the predicted junction (Fig. S7a). To test the assay, we isolated two clonal iPSC lines each with a 14.7 kb excision on chromosome 1, one matching the predicted excision repair (precise excision clone) and one with an insertion of 23 bp at the excision junction (imprecise excision clone) (Fig. S7b). The probe designed to detect the precise excision event gave equivalent results to the original ddXR assay in the precise excision clone and gave no measurable signal in the imprecise excision clone. In a mixed population of edited alleles, an average of 1.5% had a precise excision (Fig. 4a). This sample had an overall excision rate of 7.8% (Fig. 4a), suggesting that approximately 19% of all excisions underwent precise repair. We further validated this assay by performing a serial dilution of genomic DNA from the precise excision clone with genomic DNA from the imprecise excision clone and quantifying precise and total excision in each sample. The measured ratio of precise/total excision in each sample showed good agreement with the known ratio across the linear range (R^2^ > 0.98, Fig 4b).

**Figure 4:**
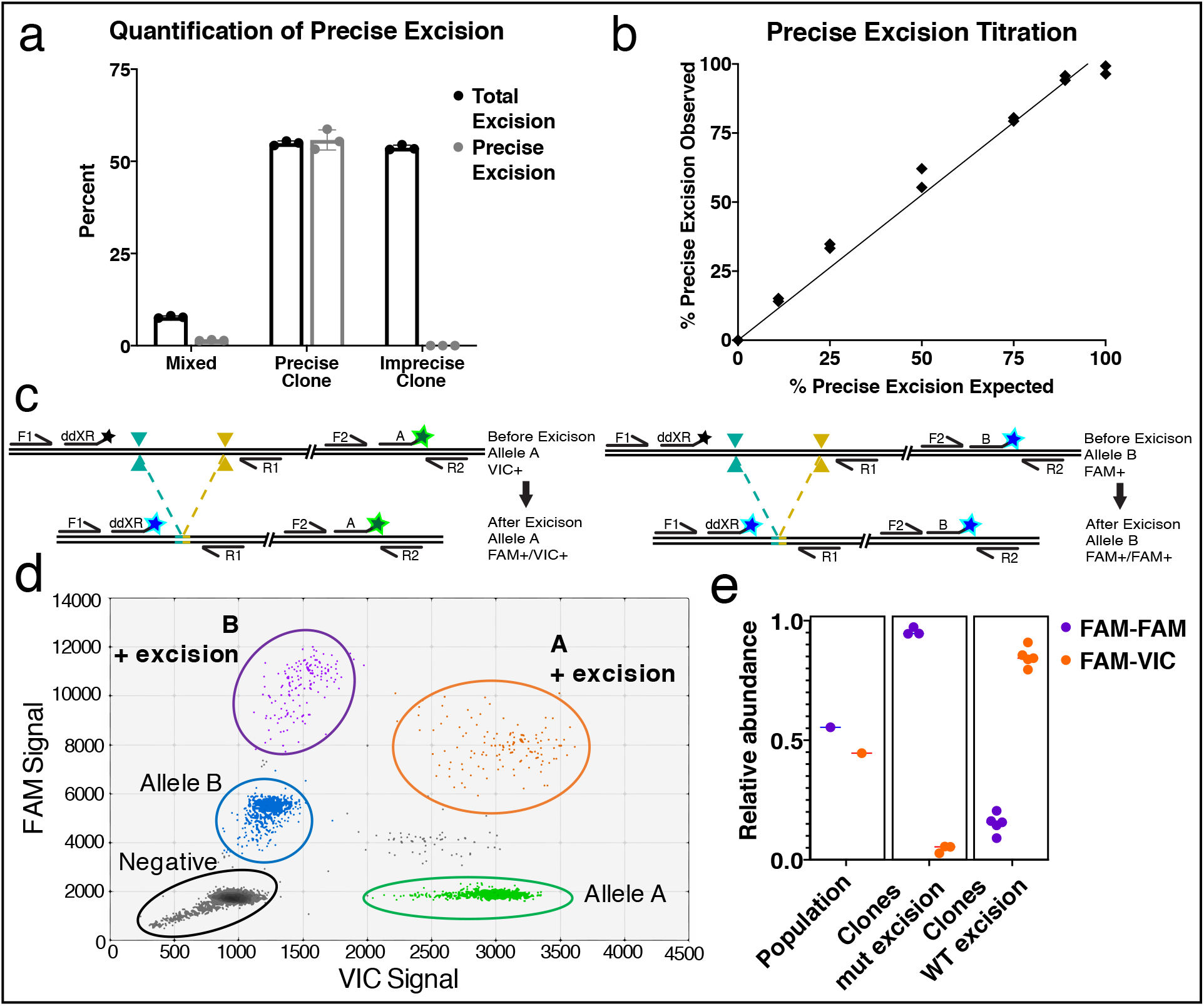
Design and validation of precise excision quantification and allele specificity assays. A) Quantification of total excision and precise excision from the mixed edited population and from edited clonal lines (Fig. S7). Values are averages of 3 replicate assays. B) Quantification of precise excisions observed by ddXR versus expected in standard samples with defined frequency of precise excision. Samples were prepared by serial dilution of genomic DNA from the clone with precise excision repair mixed with genomic DNA from the clone with imprecise excision repair. Data include two independent replicates of each dilution and are presented as the percent of excision alleles that have precise repair. R^2^ = 0.98. C) Schematic demonstrating allele-specific assay design. Allele discrimination primers (F2, R2) and probes (A = VIC / B = FAM) are designed to match a heterozygous SNP at the same locus as the excision target site, and multiplexed with ddXR primers (F1, R1) and FAM probe. Excision on allele A is expected to produce a FAM-VIC double positive signal (left), while excision on allele B is expected to produce a FAM-FAM double positive signal (right). D) Representative data from a heterozygous NEFL-N98S patient line with excision detected on both alleles. Allele discrimination probes detect unedited alleles represented by single-positive FAM and VIC populations for allele A (wild type, green) and B (mutant, blue). Excision on allele A produces FAM-VIC double positive signal (orange) while excision on allele B produces FAM-FAM double positive signal (purple). E) Quantification of excision on each allele in edited patient iPSCs and in clones. Data represent the proportion of FAM-VIC and FAM-FAM double-positive signals. Excisions occurred at similar frequency on each allele in the population, with 55.4% linked to the mutant, represented by the FAM-FAM signal (Poisson range 61.2 – 49.6). Three clones had excision on the mutant allele (mut) and five on the wild type allele, identified by predominance of either FAM-FAM or FAM-VIC signal. The FAM-FAM and FAM-VIC signals differed significantly between WT and mutant clones, and within clones the FAMFAM and FAM-VIC signal differed significantly from each other by two-way ANOVA and modified Tukey test for multiple comparisons (all p-values < 0.0001).

### ddXR detects allele-specific excisions

Allele-specific excision to delete dominant disease alleles is a promising therapeutic gene editing strategy^12^. A dominant missense mutation in *NEFL* causes a severe neuropathy that can be modeled in iPSCs to test this approach^30^. We modified ddXR to correlate excisions with specific alleles by adapting a ddPCR method previously developed for phasing SNPs on the same chromosome, using the linkage of two probes on the same molecule to produce double-positive droplets in the ddPCR readout^31^. We designed a pair of allele-discrimination probes targeting the heterozygous N98S mutation in *NEFL* (mutant-specific probe = FAM, wild type = VIC) and multiplexed them with the ddXR excision assay (Fig. 4c). We tested this assay on N98S patient-derived iPSCs transfected with a pair of RNPs not designed to be specific for either allele. In this sample, we identified FAM-FAM and FAM-VIC double positive droplets representing excision linked to either the mutant or wild type allele, respectively (Fig. 4d). As expected, we observed similar rates of editing at each allele in the population of edited cells (Fig. 4e). We next isolated iPSC clones from the edited population and used the allele-specific ddXR assay to identify clones in which excision occurred on a specific allele. Clones with heterozygous excision linked to wither mutant or wild type allele were identified by the presence of double-positive droplets, either FAM-FAM or FAM-VIC, respectively (Fig. 4e and Fig. S8). A single homozygous clone with excision on both alleles was identified by presence of both double-positive signals (Fig. S8d). These data confirm that heterozygous SNPs adjacent to the site of an excision can be incorporated into the ddXR design to determine whether an excision is specific for a particular allele.

## Discussion

Here, we show that ddXR provides an accurate and length-independent method for quantifying the frequency of inversions and excisions in genome editing experiments. This is especially important for applications that require deletions larger than a few hundred base pairs. While excision length is limited in studies that rely on NGS for quantification^8,16,17^, we are able to quantify excisions up to 172 kb and have not yet reached an upper limit to the length of excision or inversion detectable by ddXR. We compared ddXR to other methods, including clone genotyping and FISH, which produced comparable but lower throughput results. We further showed that ddXR outperforms a LOS ddPCR assay, displaying higher precision across a wide range of editing efficiencies. Therefore, ddXR fills a need in the field for a flexible and rapid method to quantify excisions and inversions. We performed all of the assays and validation in well-characterized, euploid human iPSC lines, making our results maximally relevant to human gene editing.

Our modifications for the quantification of precise repair and allele specificity demonstrate the versatility of this assay (Table 1). The ability to design and carry out assays for such varied editing outcomes also points to the consistency of the assay. Though there are some design constraints for the sequences of the probes and primers, we have not yet encountered an excision for which we were unable to design an effective assay. However, larger than expected deletions that are suggested by our FISH results (Fig. S6) and others’ reports^32–34^, would be missed. Although a single ddXR assay cannot measure all possible excision sizes and rearrangements, further adjustments, similar to the inversion assay, can be easily designed to quantify specific rearrangements of interest once they are defined. It is likely that other modifications will further expand the scope of ddXR. For example, a current limitation of the assay is that we cannot distinguish between heterozygous and homozygous excisions or inversions in a polyclonal sample, as all of the alleles are assayed in bulk. Single-cell modifications of the protocol would be a welcome future improvement.

**Table 1:** Summary of ddXR assays and modifications.

| Assay | Primer Design | Probe design | Control | Notes |
| --- | --- | --- | --- | --- |
| **ddXR - Excision** | Forward and reverse primers flank excision junction | Flanking either 5' or 3' target site | Genomic reference (e.g. RPP30) | Add restriction digest for excisions < 200 bp |
| **ddXR - Inversion** | Two forward primers (2<sup>nd</sup> replaces reverse excision primer, internal to cut sites) | Same as excision | Genomic reference (e.g. RPP30) |  |
| **Precision ddXR** | Same as excision/inversion | Spans predicted repair junction | Genomic reference *or* general ddXR | Normalize to RPP30 for absolute *or* ddXR for relative |
| **Allele - specific ddXR** | Same as excision/inversion | Same as excision/inversion | Allele-discrimination to nearby SNP | Double-positive droplets identify linkage to SNP |

This assay provides a robust tool for evaluating the determining factors that promote excision and inversion. Based on the varying frequencies we observe across a wide range of linear DNA length, it is likely that many factors affect excision and inversion formation. In contrast to a previous study^3^, we did not observe a strong decrease in excision frequency with increasing length and observed no correlation between inversion frequency and length. The previous study reported nearly undetectable frequency of excisions larger than approximately 20kb in length, as measured by clone genotyping in an immortalized murine cell line, while we were able to detect both excisions and inversions up to 172 kb in length at surprisingly high frequencies. Differences in the species and type of cell lines could partially explain the differing outcomes of our experiments. It is also probable that some very large deletions compromise cell viability and/or proliferation, which would prevent the cells from establishing clonal populations. Conversely, our method requires only a small amount of DNA and can be performed at any time point after editing, without expansion of the edited populations. It should therefore be possible to study editing outcomes even in non-proliferative cells, such as primary cells and various differentiated cells derived from iPSCs.

Rigorous studies of the repair of single DNA cleavages induced by CRISPR-Cas have begun to reveal predictable patterns that can be used to promote a specific repair outcome^35,36^. Until now, the lack of suitable methods has slowed the discovery of similar rules for excision-linked DNA repair. Our data contradict the hypothesis that linear distance is the primary factor driving the frequency of excision or inversion and ddXR enables systematic interrogation of other potential determinative variables including different nuclease enzymes, indel activity, PAM orientation, chromatin state and 3D DNA structure. The additional ability to measure precise repair and allele-specificity is particularly relevant to disease modeling, where targeted deletion of specific sequences allows for the detailed study of structure-function relationships, and for therapeutic editing. An exciting application of allele-specific excision is therapeutic inactivation of dominant disease alleles. In this regard, we focused our studies on three candidate loci containing genes that cause severe dominant inherited neuropathy (*HSPB1, MFN2 and NEFL*). The methods described here represent an important advance toward development of effective therapeutic editing strategies for these and other severe diseases.

## Acknowledgments

We thank Mario Saporta for providing iPSC lines from neuropathy patients, the Gladstone Stem Cell Core and Cell Line Genetics for their services, and A. May, G. Cirolia, B. Wienert, J. Perez-Bermejo and C. Clelland for helpful discussions and critical review of the manuscript.

## Author contributions

L.M.J., H.L.W. and B.R.C. conceived the project and designed the experiments. K.G. conceived of the allele-specific assay design. Y.M. and G.T. conceived of the precision assay design and performed initial validation. H.L.W. and C.M.F. performed the experiments. H.L.W. and L.M.J. conducted data analysis and wrote the manuscript with input and assistance from all authors.

## Methods

### iPSC culture and CRISPR editing

WTC^37^ and patient cell lines, MN-P1^30^, NF-P1^30^, HB-P1^18^ are all previously published and characterized. All iPSCs were cultured in Stemfit (Ajinomoto) on plates coated with matrigel (Corning) at 37 °C, 5% CO_2_ and 85% humidity. gRNAs were designed using CRISPOR^38^. Ribonucleoproteins (RNP) were prepared by complexing 104 pmol of each guide RNA (IDT) with 52 pmol of spCas9 protein (MacroLab) prior to transfection. Cells were transfected with RNPs using the Lonza 4D-Nucleofector X unit with pulse code DS138. After nucleofection, cells were cultured in media with Rock Inhibitor Y-27632 (SelleckChem). Edited samples for ddXR assay were harvested 2 to 4 days post nucleofection without passaging. Clonal cell lines underwent two rounds of manual clone picking followed by expansion until an adequate number of cells for both cryopreservation and DNA extraction was obtained. DNA was extracted from non-clonal samples using the DNeasy Kit (Qiagen). DNA for clone screens was extracted by QuickExtract (Lucigen) or ethanol precipitation.

### ddPCR assays

ddPCR primer/ probe pairs were designed with the Primer Express software, using the MGB Quantification option. FAM-conjugated probes were designed to match sequence on the outside of one of the excision gRNAs, approximately 20 bp away (Fig. 1a). One primer was placed just outside the probe from the cut site, ideally 40 bp away from the cut site. The second primer was placed just outside the second cut site. To detect inversions, the second primer was designed within the cut region pointing outwards adjacent to the second cut site. Ideally, amplicons should be between 100 and 150 bp. For repetitive or GC rich regions, we designed primer sets that produced amplicons up to 200 bp. Probes to detect precise excisions were designed to overlap the predicted junction, with the breakpoint in the center of the probe (Fig.S7a). The SNP probes used for allele-specific ddXR were designed using the Primer Express MGB Allele Discrimination option. A pair of VIC/FAM probes were designed for a SNP near, but not within, the excision. The SNP should be chosen as close as possible to the excision target site to minimize disruption to linkage caused by DNA shearing (Fig. 4c). All reactions, except for allele-specific ddXR, include a HEX-conjugated probe and primers to the RPP30 gene (Bio-Rad Laboratories) on chromosome 10, which encodes ribonuclease P/MRP subunit p30 and serves as an internal control to normalize the frequency of the gene editing outcome in question. A probe to any reference gene present in two copies can be used in place of RPP30. Sequences of gRNAs, PCR primers and probes are provided in Tables S2–S6.

All 25 uL ddPCR reactions were composed of 12.5 uL Supermix for Probes (no dUTP) (Bio-Rad Laboratories), 1.25 uL 20X reference assay, 50 ng DNA, 1.25 20X FAM assay for the target edit and water to 25 uL. Each 20X target probe mixture was made of 18 uM forward and reverse primer each and 5uM target probe. Droplets were generated using 20 uL reaction mixture and 70 uL oil with the QX200 Droplet Generator (Bio-Rad Laboratories). Droplets were transferred to a 96-well PCR plate, sealed and run on a C1000 Thermal Cycler with a deep-well block (Bio-Rad Laboratories). For samples with the 91 bp deletion, a restriction digest was performed prior to PCR amplification. 500 ng of DNA was treated with FastDigest BstXI in FastDigest Buffer (Thermo Scientific). Restriction digests were incubated at 37 °C for 60 min followed by heat inactivation at 65 °C for 20 min.

All ddPCR reactions were run under the following thermal cycling conditions: 1) 95 °C for 10 min; 2) 94 °C for 30 s; 3) 58 °C for 1 min; 4) steps 2 and 3 repeated 39 times; 5) 98 °C for 10 min.

All ddPCR runs were analyzed using the Bio-Rad QuantaSoft Pro Software. For inversion and excision rates, the value listed as “ratio” (excision or inversion to RPP30) in QuantaSoft was used. For allele-specific ddXR, the ratio of VIC+/VIC+ events and FAM+/VIC+ events were used to calculate the ratio of edited alleles.

### Clone genotyping

All clones were genotyped by PCR using primers placed outside the excision. PCR products were run on a 1% agarose gel to check for the presence of an excision band. For short excisions, the larger, unedited amplicon may be visible. For inversions, a primer 5’ to each gRNA site was used to detect inversions. The product is only amplified after the inversion occurs as it reverses to orientation of the second primer. All primer pairs were tested on unedited DNA control to confirm there was no PCR band prior to excision or inversion.

### Generation of genomic DNA standards

DNA from a pure, edited clone was mixed with unedited DNA of the same parent cell line. A serial dilution was performed, each with a total volume of 30 uL. The amount of edited DNA was decreased by a factor of four with each dilution with additional data points at 1.56, 0.39 and 0.09 (1:2 dilutions) for better resolution at lower concentrations. All dilutions were made and quantified in technical duplicates. For the 4.09 kb excision, the same DNA dilutions were used for standard curves for both the GOS and LOS assays. R^2^ and RMSE values were calculated using Prism.

#### Fluorescent in-situ hybridization

FISH was performed by Cell Line Genetics Inc. (CLG) in Madison WI. A cryopreserved vial of unedited WTC cells and WTC cells edited with guides chr1:2 and chr1:4 were sent to CLG. CLG performed FISH on 200 cells for each sample using probe 1p36.22 (BAC clone RP11-1005H15 chr1:11,821,652-11,999,215) to detect the deleted region and probe 1p36.13 (BAC clone RP11-1062E1 chr1:17,114,168-17,348,163) as a reference. Cells from the same sample and passage were used to extract DNA for ddXR quantification.

## Supplemental Figures

**Figure S1:**
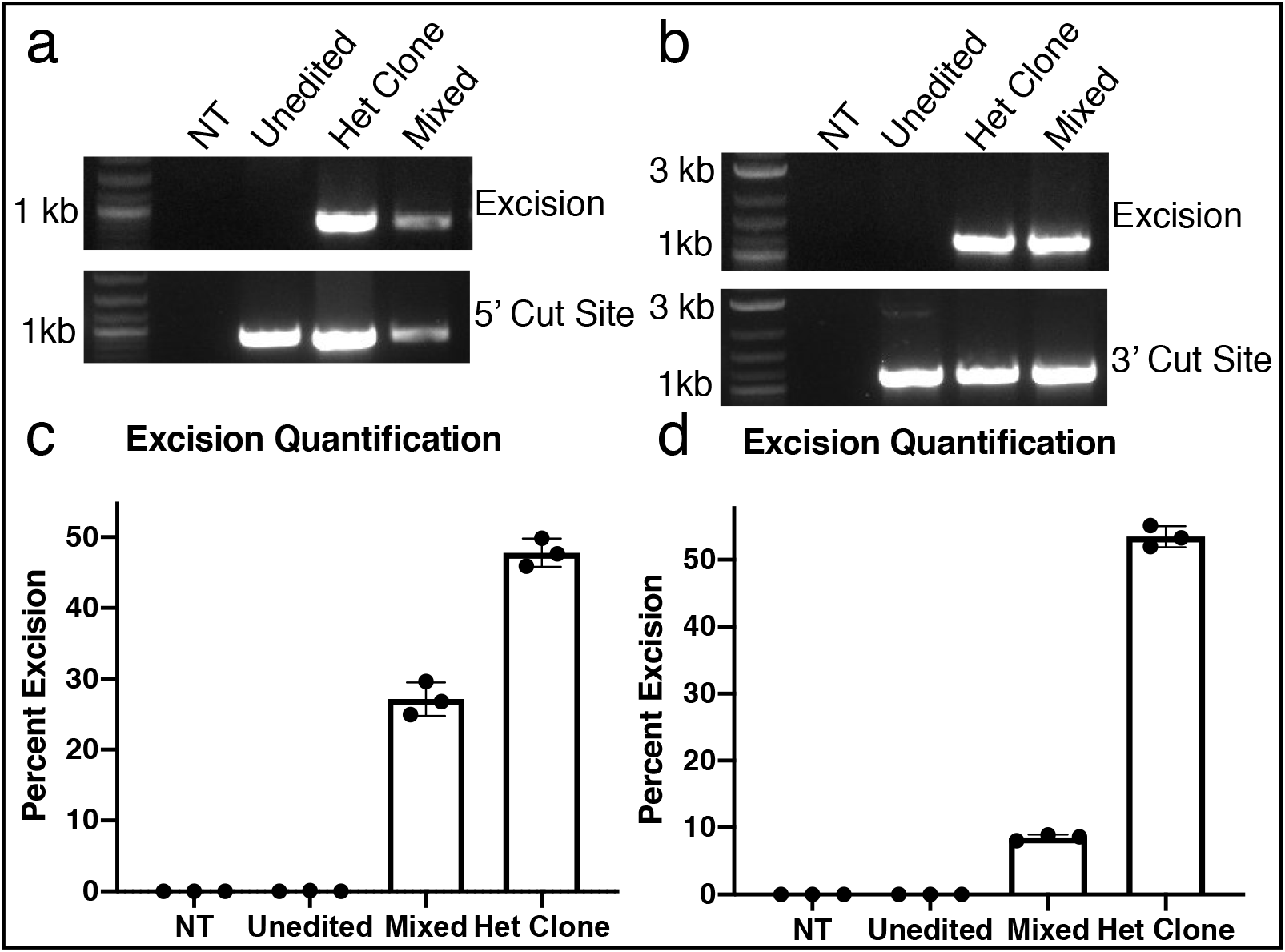
Validation of ddXR for excision quantification at 2 additional loci using clonal iPSC lines. A,B) Validation of clonal line with heterozygous 1.7 kb excision on chromosome 7 (A) and heterozygous 14 kb excision on chromosome 1 (B). PCR spanning excision produced expected bands in the mixed sample and clone. Control PCR spanning 5’ or 3’ cut site showed bands in the unedited, clone and mixed sample. C,D) ddXR quantification of 1.7 kb (C) and 14 kb (D) excision in negative controls (no DNA and unedited) and heterozygous clone and mixed sample. Values are averages of 3 replicates.

**Figure S2:**
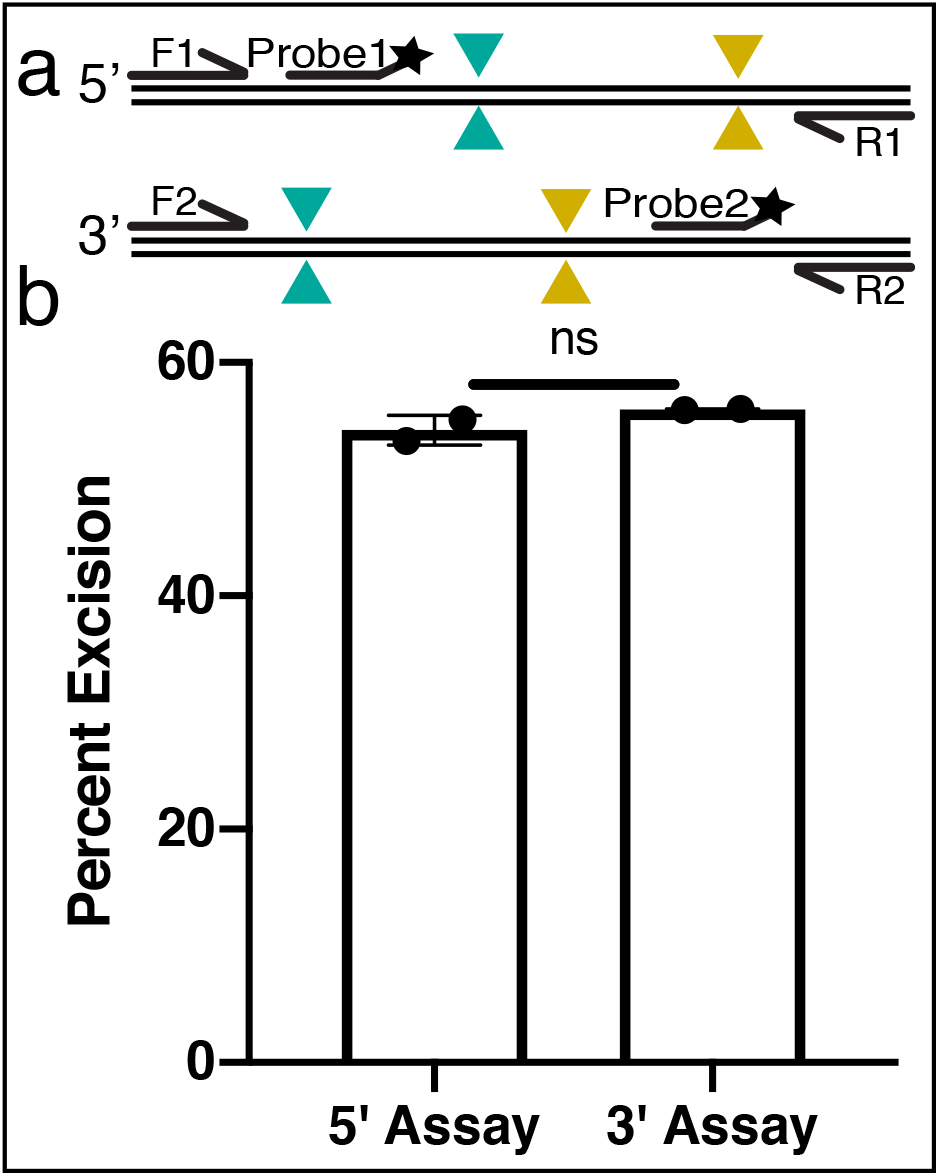
Evaluation of the directionality of the ddXR assay. A) Diagram of 5’ and 3’ version of assay. B) Quantification of frequency of 14 kb excision in a clonal iPSC line using 5’ and 3’ assays. Values are averages of two replicates. p = 0.2879.

**Figure S3:**
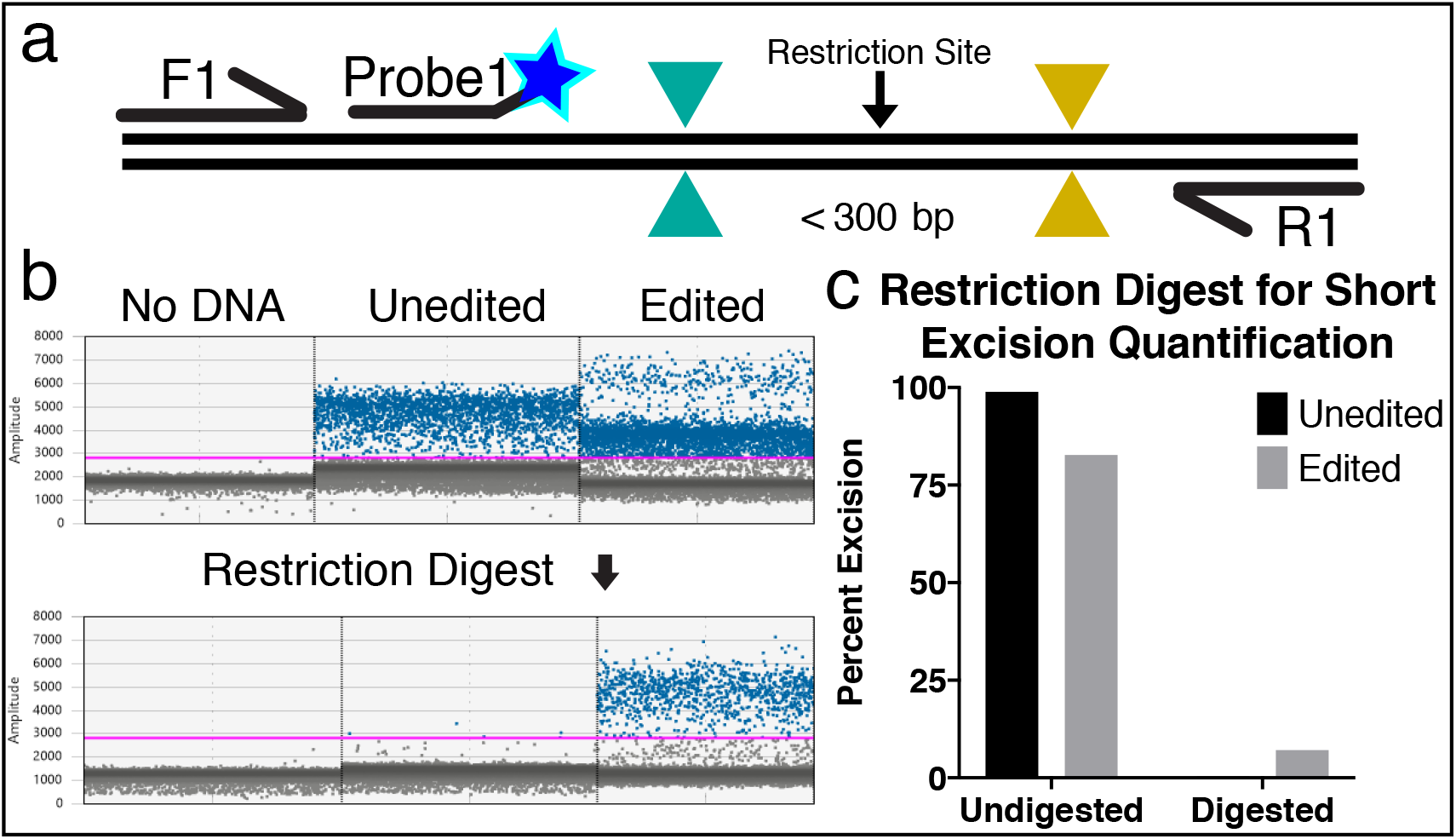
Restriction digest eliminates false-positive signal when quantifying short excisions. A) Schematic of restriction digest recommended for short excision samples. B) Representative 1D data plots from ddXR quantification of a 91 bp excision on chromosome 1 with and without restriction digest with BSTXI FD enzyme. C) Percentage of excisions detected in unedited and edited DNA with and without restriction digest, using the gating thresholds indicated by the pink bar in (B). Restriction digest effectively removed false positive signal caused by efficient amplification of non-excised alleles. This simple modification can be performed on any short excision using a restriction enzyme recognition sequence present within the excised region, but not elsewhere within the expected amplicon. Short inversions do not present the same challenge since the orientation of the primers does not allow amplification of unedited alleles, regardless of length.

**Figure S4:**
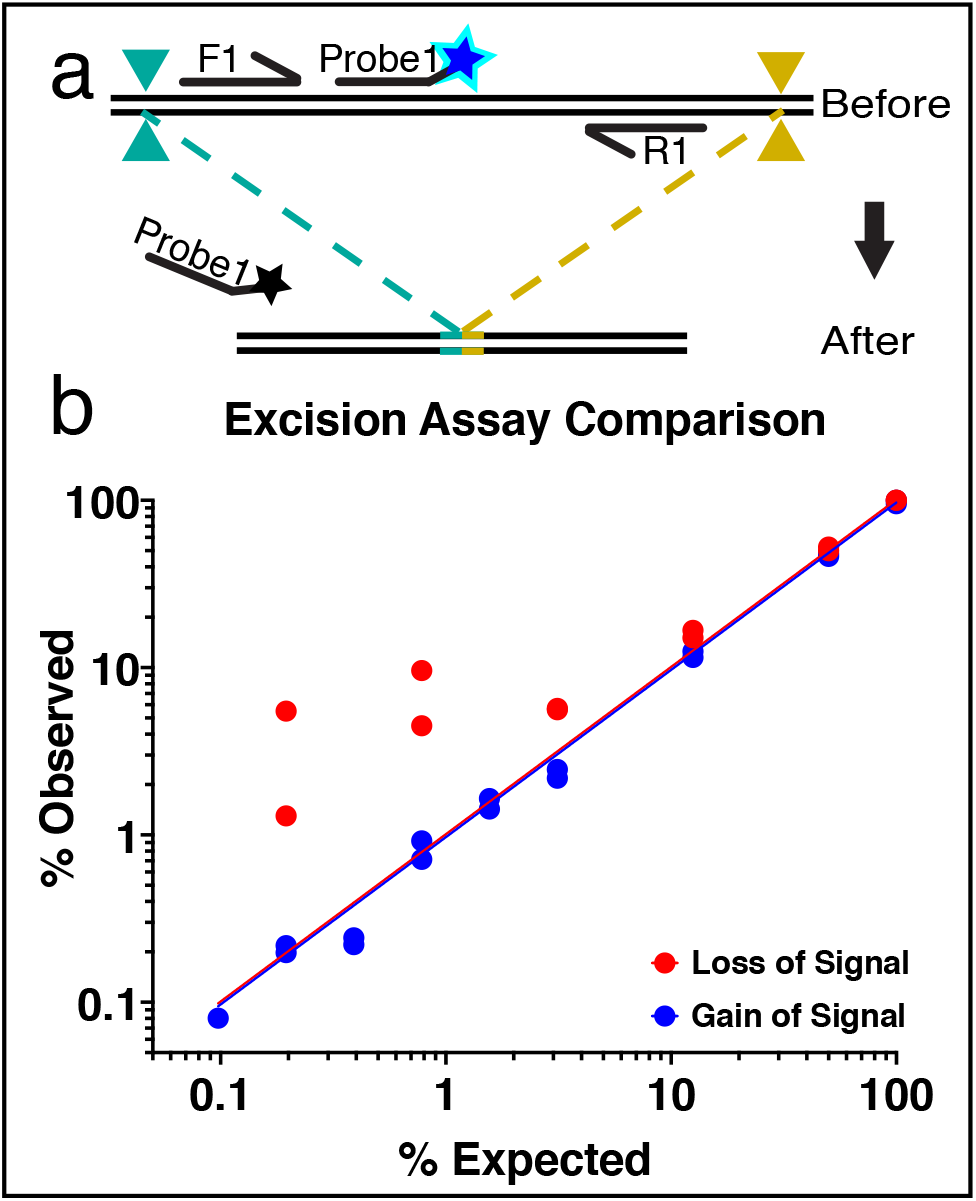
Excision quantification by loss of signal (LOS) ddPCR assay. A) Schematic of LOS ddPCR assay for excision quantification. B) Comparison of ddPCR quantification of standard samples with defined frequency of a 4.09 kb excision by LOS (red) and ddXR (blue) assays. Samples are the same as those shown in Fig 2e. RMSE for LOS = 4.161. RMSE for ddXR = 0.9924.

**Figure S5:**
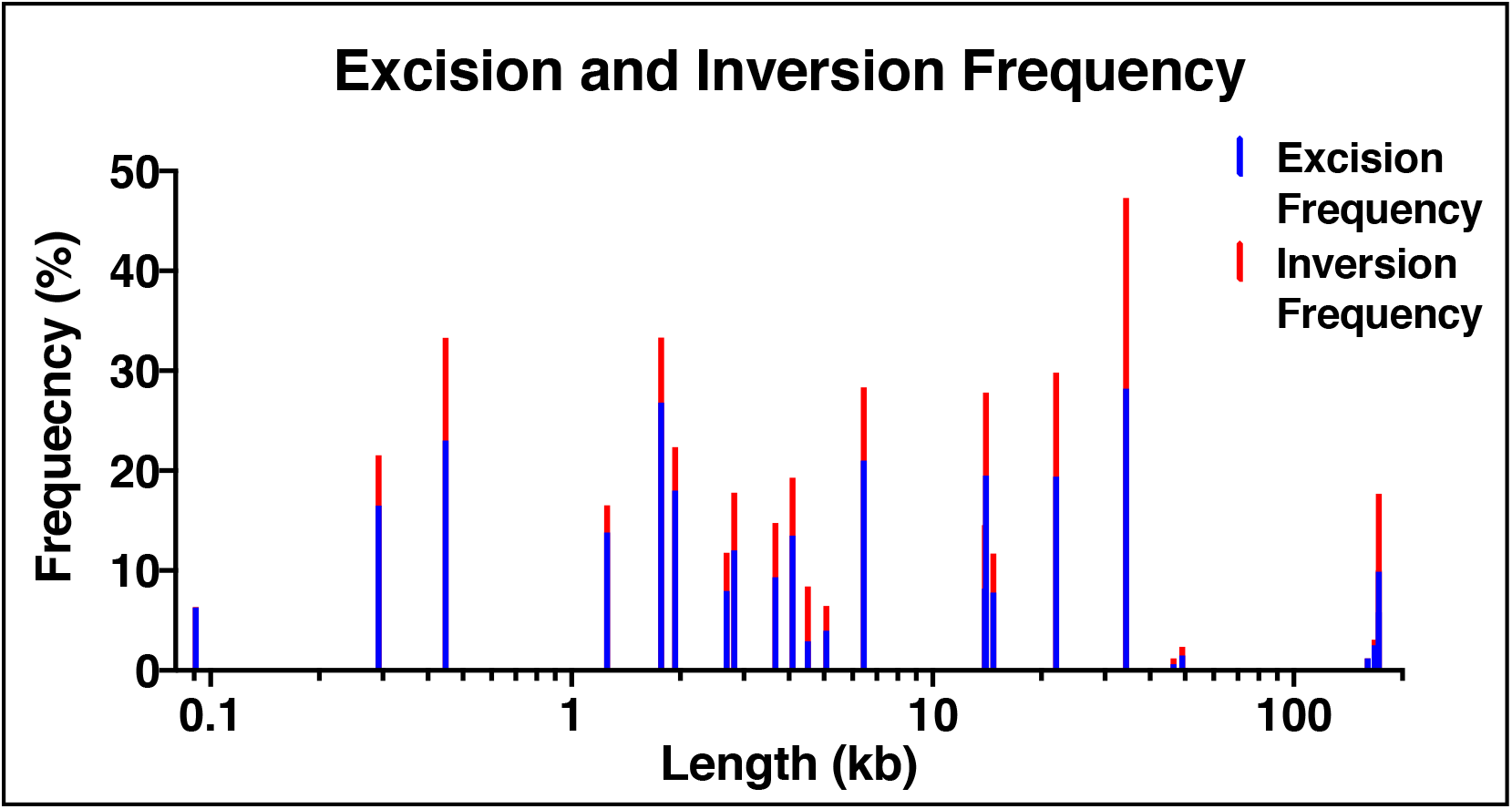
Quantification of excision and inversion frequency across a range of linear chromosomal distances. Stacked excision (blue) and inversion (red) rates of paired CRISPR-Cas9 ranging from 91 bp to 172 kb apart. Data are the same as presented in Fig. 3 and represent combination of experiments targeting three distinct loci each on distinct chromosomes.

**Figure S6:**
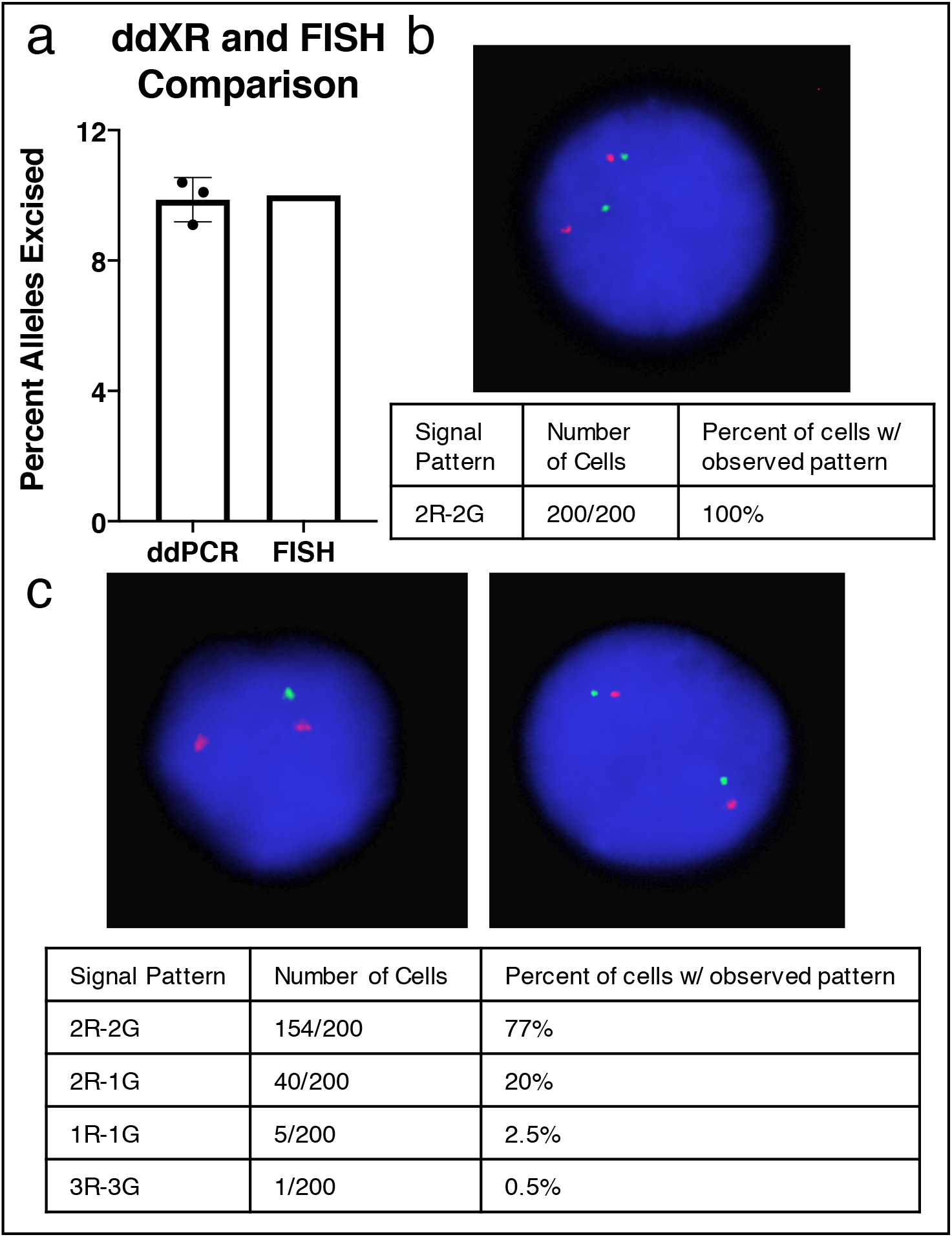
Validation of large mixed population excision rate by FISH. A) Comparison of quantification of 172 kb excision by ddXR and FISH. ddXR value is an average of three replicate assays. B,C) FISH result table and representative images from unedited (B) and edited (C) samples. Red Probe = Control. Green Probe= experimental (within excision). Cells with both alleles intact have 2 red and 2 green signals, cells with 1 green and 2 red represent a heterozygous excision event. C) Edited sample includes five cells with a single copy of both reference and experimental probes. While these could be FISH artifacts, it is possible that these cells had a larger than expected (>5 Mb) excision that ablated both the target sequence and the sequence recognized by the reference probe.

**Figure S7:**
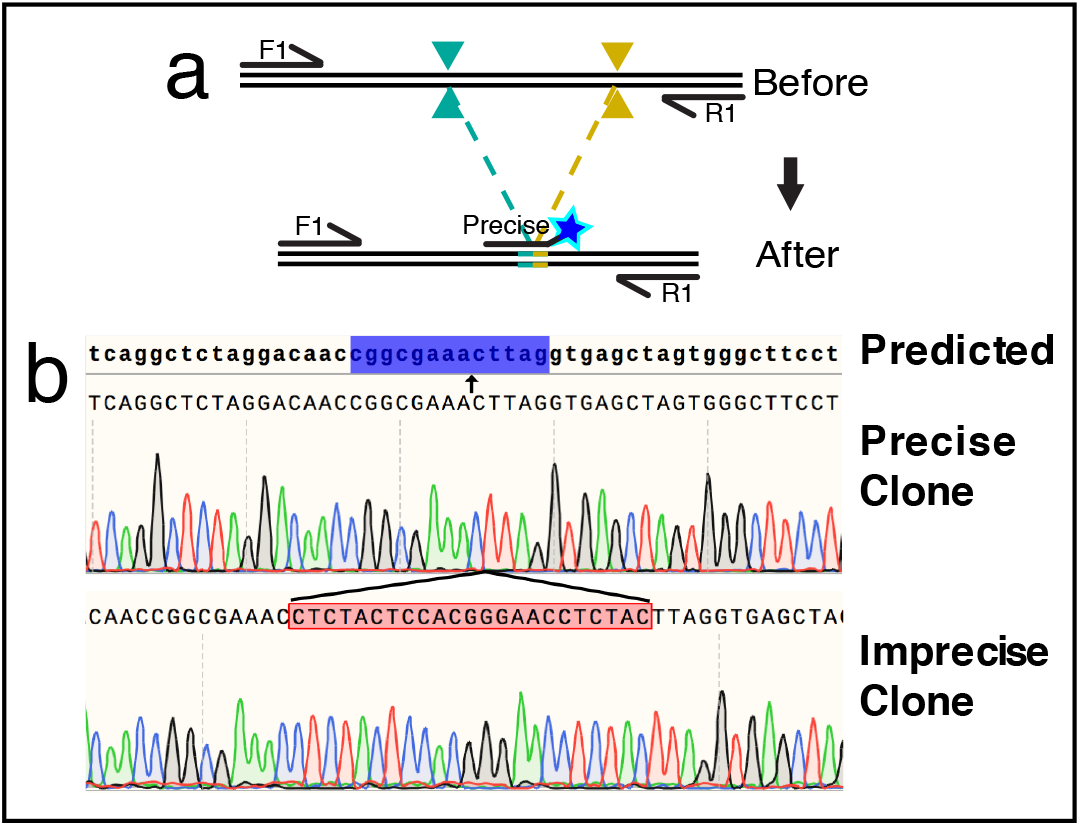
Precise excision assay and characterization of precise and imprecise excision clones. A) Schematic of precise excision assay with probe spanning excision repair junction. B) Sanger sequencing data from clonal iPSC lines with a 14kb excision on chromosome 1, one with a precise repair of the predicted excision junction (precise clone) and one with a 23bp insertion at the excision junction (imprecise clone). Probe designed to detect precise repair junction is highlighted in blue, with arrow indicating the junction between cut sites.

**Figure S8:**
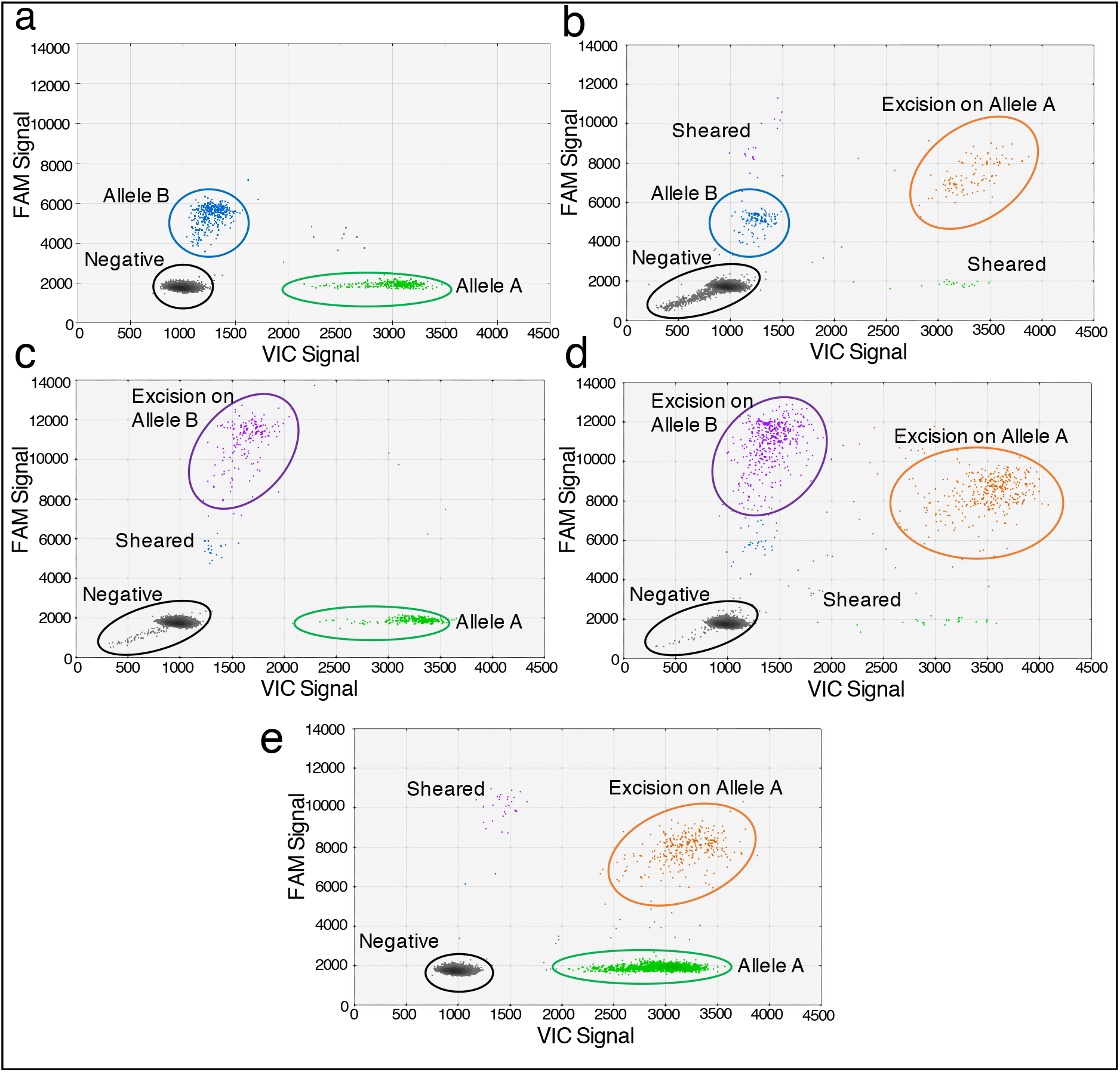
Validation of allele specific ddXR on unedited, clonal, polyclonal lines. A-E) Additional representative 2D data plots from unedited control (A), clonal line with excision on allele A (B), clonal line with excision on allele B (C), clonal line with homozygous excision (D) and edited population of cells homozygous for allele A (E). Allele discrimination probes detect unedited alleles represented by single-positive FAM and VIC populations for allele A (wild type, green) and B (mutant, blue). Excision on allele A produces FAM-VIC double positive population (orange) while excision on allele B produces FAM-FAM double positive (purple). The small number of single-positive droplets in (B-E) are produced by shearing of DNA that disrupts the linkage between the SNP and excision site in a subset of templates. The source of this population is most apparent in (E), where the purple population is present, despite the cell line lacking the mutant allele B. The ddXR excision probe produces a FAM signal that partially overlaps with the FAM-FAM double-positive signal. This single-positive FAM signal produced by shearing explains the relatively higher FAM-FAM measured signal in clones with excision on the wild-type allele. Decreasing distance between the SNP and the excision site is expected to minimize this shearing phenomenon.

## Supplemental Tables

**Table S1:**
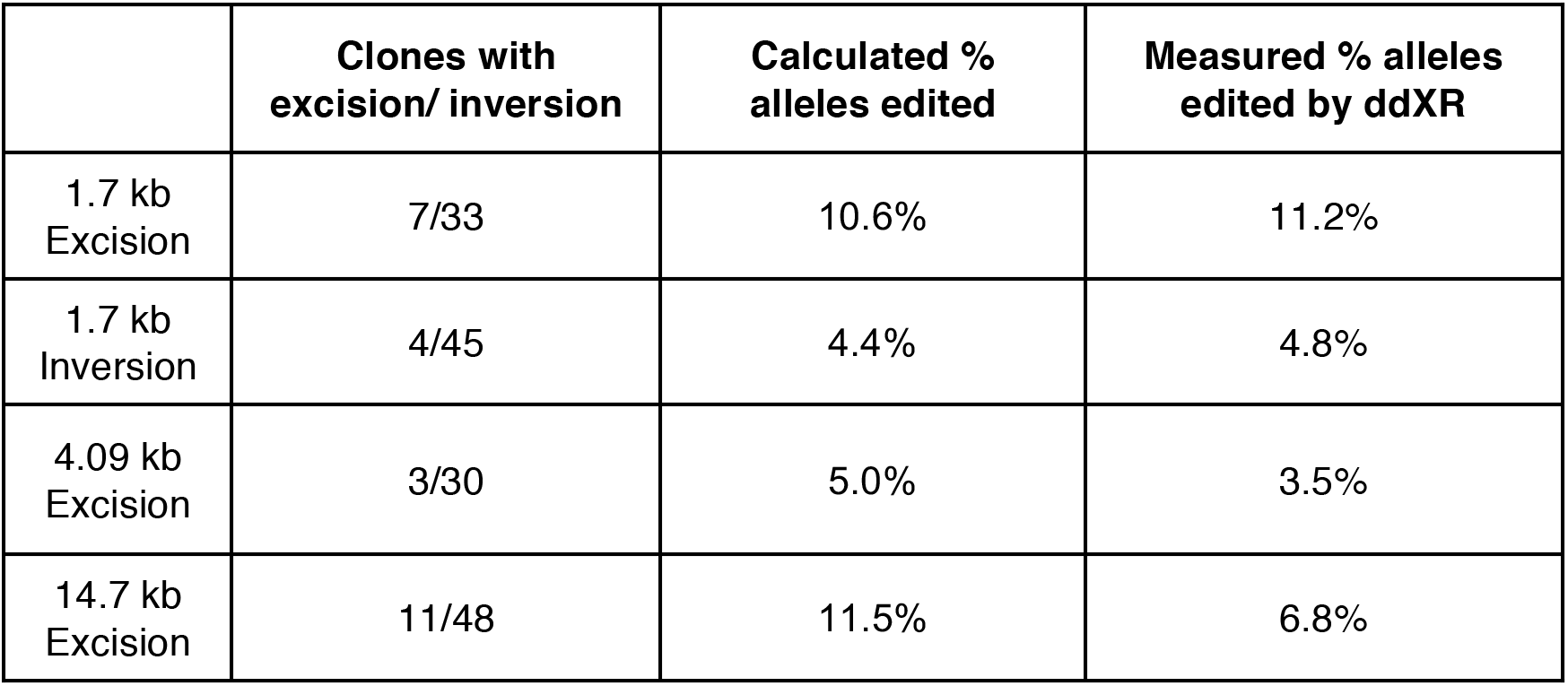
Comparison of excision/ inversion rates determined by genotyping of clones and by ddXR.

**Table S2:**
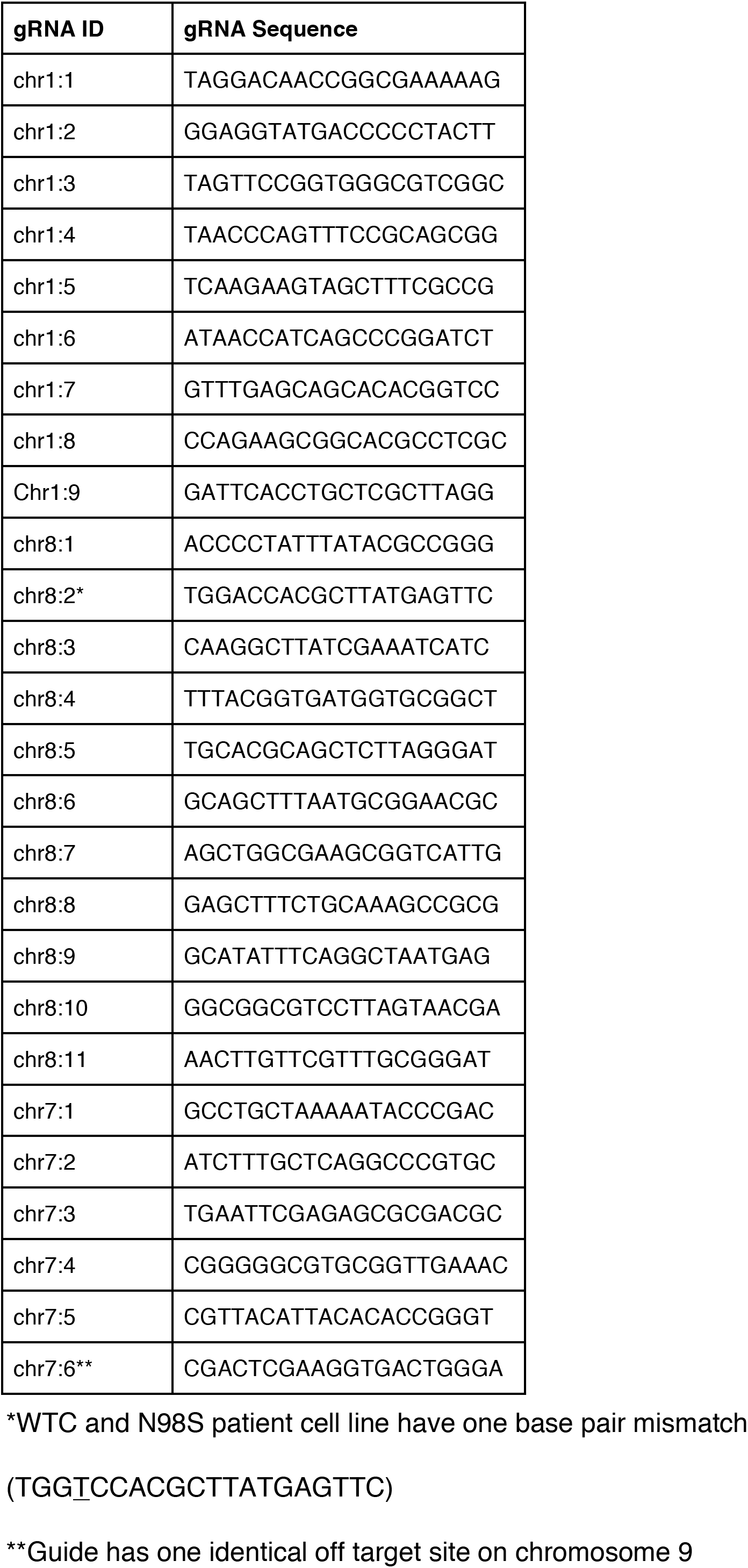
List of gRNAs used in excisions. gRNAs are identified by the chromosome they align to.

**Table S3:** gRNA pairs and cut lengths. 5’ guide is listed first.

| **gRNA Pairs** | **Length (bp)** |
| --- | --- |
| chr1:1/chr1:2 | 14,698 |
| chr1:1/chr1:3 | 91 |
| chr1:1/chr1:5 | 6,443 |
| chr1:1/chr1:6 | 14,732 |
| chr1:1/chr1:7 | 21,986 |
| chr1:4/chr1:2 | 172,082 |
| chr1:8/chr1:2 | 160,232 |
| chr1:9/chr1:2 | 171,946 |
| chr1:8/chr1:7 | 167,485 |
| chr8:1/chr8:2 | 4,087 |
| chr8:3/chr8:2 | 5,078 |
| chr8:4/chr8:2 | 4,511 |
| chr8:5/chr8:2 | 2,821 |
| chr8:6/chr8:2 | 2,689 |
| chr8:7/chr8:2 | 3,667 |
| chr8:9/chr8:2 | 46,433 |
| chr8:10/chr8:2 | 49,135 |
| chr8:1/chr8:8 | 14,017 |
| chr8:1/chr8:11 | 13,960 |
| chr7:1/chr7:2 | 1,750 |
| chr7:3/chr7:2 | 1934 |
| chr7:4/chr7:2 | 1254 |
| chr7:5/chr7:2 | 448 |
| chr7:6/chr7:2 | 292 |

**Table S4:**
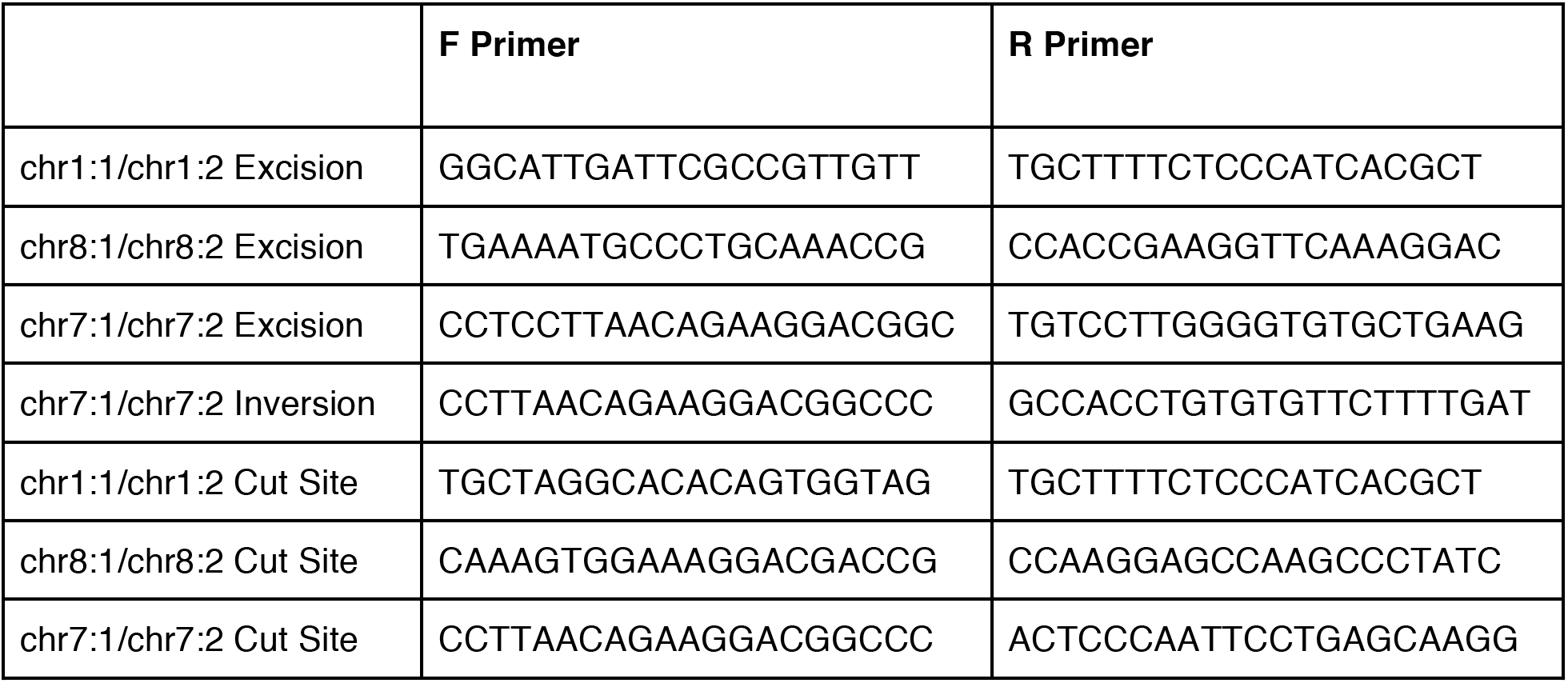
Standard PCR primers used to genotype clones in Fig.2a,b, S1a,b and Table S1.

**Table S5:**
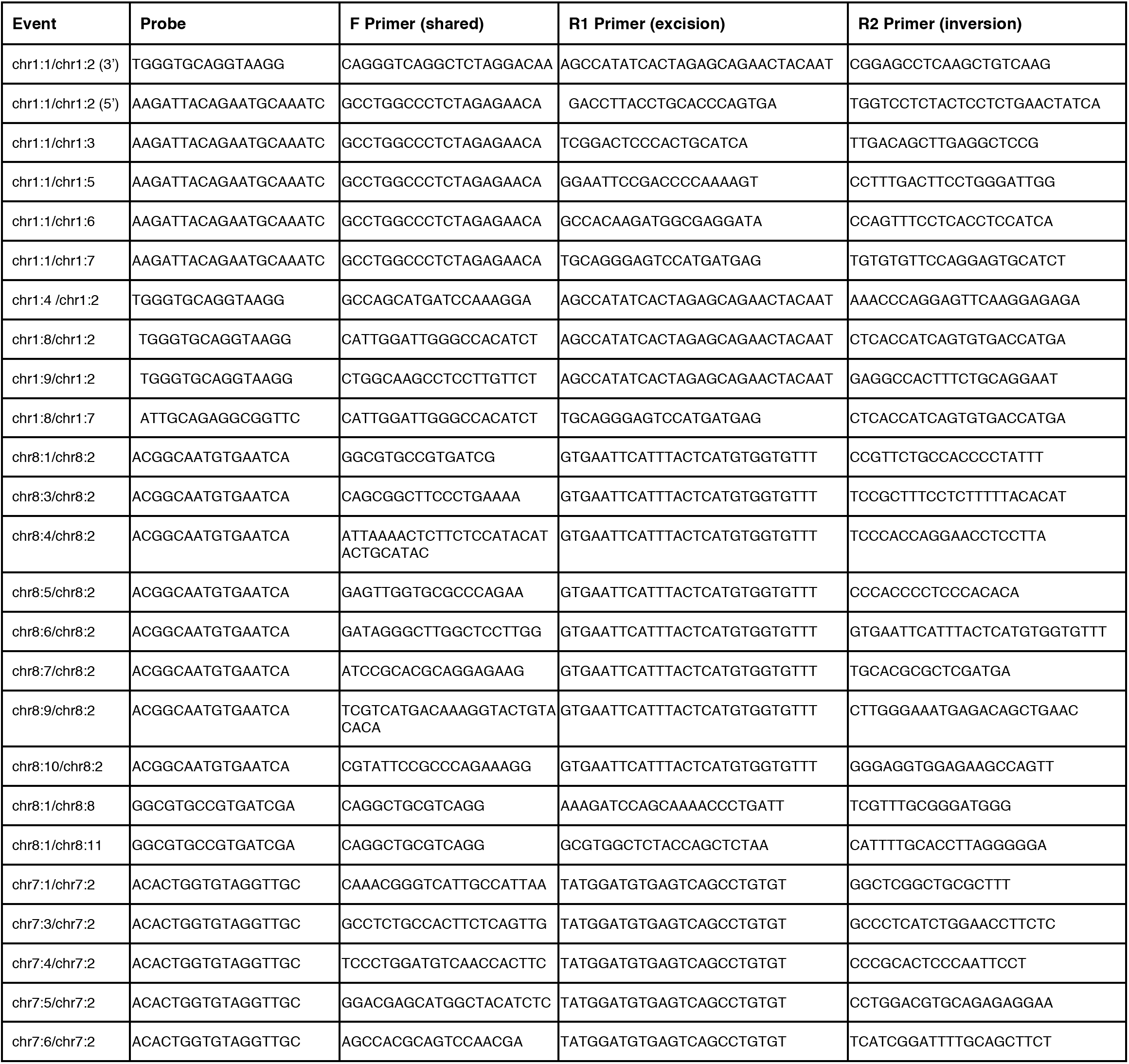
Probes and primers for all ddXR assays.

**Table S6:**
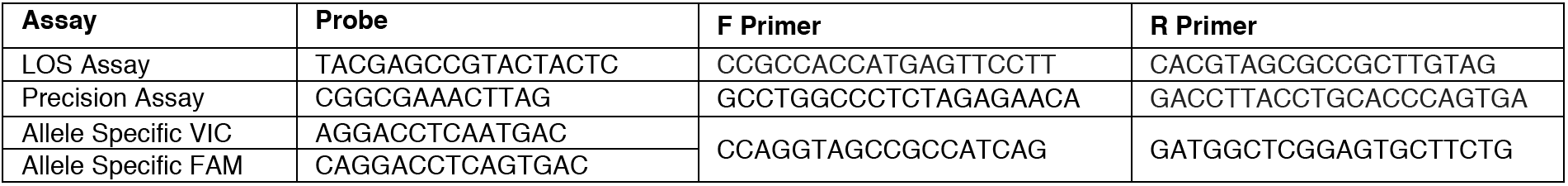
Additional probes and primers used.

